# Light at the end of the tunnel: FRAP assay reveals that plant vacuoles start as a tubular network

**DOI:** 10.1101/2021.05.13.444058

**Authors:** Elena A. Minina, David Scheuring, Jana Askani, Falco Krueger, Karin Schumacher

## Abstract

Plant vacuoles play key roles in cellular homeostasis performing catabolic and storage functions, regulating pH and ion balance. The essential role of vacuoles for plant cell viability makes them a notoriously difficult subject to study impeding reaching the consensus on the mechanism of vacuolar establishment and the source of membrane material for it. Our previous suggestion of endoplasmic reticulum being the main membrane contributor for the tubular network of young vacuoles was recently challenged in a study proposing that young plant vacuoles comprise a set of individual vesicles that are formed *de novo* via homotypic fusion of multivesicular bodies (MVBs).

To resolve these seemingly contradictory observations we have carefully revaluated both hypotheses. Here we provide a systematic overview of successive vacuolar biogenesis stages in Arabidopsis root, starting from the youngest cells proximate to the quiescent center. We validate our previous conclusions by demonstrating that the vacuolar dye BCECF is fully suitable for studying the organelle’s morphology and provide 3D models of vacuoles at all developmental stages. Furthermore, we established a customized FRAP assay and proved that even at the earliest stages of biogenesis, vacuoles comprise a connected network. Finally, we summarized the new and pre-existing evidence substantiating that vacuolar structures cannot originate solely from MVBs.

## Main

Vacuoles are the largest organelles of plant cells, they play the key role in cellular homeostasis performing catabolic and storage functions, regulating pH and ion balance and are vital for plant cell adaptability to environmental changes^1–5^. These organelles are highly dynamic and vary greatly in size, morphology and content, depending on plant species, cell type, developmental stage and environmental conditions^4,5^. Plant vacuoles typically undergo massive structural changes during cell differentiation and growth, starting as relatively small organelles in meristematic cells and occupying up to 90% of the cellular volume in differentiated cells^3,6^. It is intriguing how such vast reorganization of the organelle is synchronized with its vital role in cellular homeostasis and responses to environmental stress stimuli. It is also an exciting question, what is the source of the vacuolar membrane (tonoplast) that supports the massive reorganization and growth of the organelle^3,7,8^.

The presence of the vacuoles in all plant cells implies that these organelles are likely to be inherited from cell to cell, but does not exclude that a subpopulation of the vacuolar structures might be formed *de novo*.

Understanding the mechanisms underpinning vacuolar biogenesis is crucial for elucidating the role of these organelles in plant physiology. However, obtaining the required knowledge has been a challenging task due to the pleiotropic effects of mutations impacting vacuolar biogenesis^9–11^ and embryo lethality of the mutants with impaired vacuolar establishment^12^. For this reason, the over a century old debate about the origin of plant vacuoles is still ongoing^7,8,13,14^.

So far it has been suggested that plant vacuoles might originate from plastid-like structures^14^, or from Golgi-associated endoplasmic reticulum (ER) that undergoes autophagy-dependent modifications^15^, or are established with participation of ER-derived provacuoles^7^. The recent work by Cui et al.^16^ proposed an intriguing hypothesis, according to which plant vacuoles start as small vacuoles (SVs) that are *de novo* produced in young meristematic cells *via* homotypic fusion of multivesicular bodies (MVBs). However, the existing body of evidence strongly suggests that homotypic fusion of MVBs would not produce fully functional vacuolar structures.

For instance, in agreement with the pre-existing data, Cui et al. demonstrated that vacuolar structures and MVBs are decorated with two different sets of membrane proteins. This observation alone already indicates that formation of the vacuolar membrane is unlikely to rely solely on MVBs. The process plausibly requires an additional membrane source that would deliver the characteristic transmembrane proteins. Indeed, as we have demonstrated previously, the transmembrane subunits of two highly abundant tonoplast protein complexes present on all vacuolar membranes, the vacuolar H^+^-ATPase and the vacuolar H^+^-PPase are delivered to the vacuole via an MVB-independent trafficking route^7^.

Post-Golgi trafficking in general and MVBs in particular undoubtedly play an important role in plant vacuole establishment. MVBs are essential for the trafficking of proteins from ER and plasma membrane to the vacuole.^17^ During this process, intraluminal vesicles (ILVs) of MVBs accumulate cargo sorted for delivery to the vacuole and are released into the vacuolar lumen upon MVBs fusion with the tonoplast^17^. The presence of ILVs in the vacuolar lumen reported by Cui et al. is thus a well-described phenotype merely indicative of a functioning endomembrane trafficking towards the vacuole and does not speak for vacuoles originating from MVBs^16^. The mechanism regulating cargo sorting, formation of ILVs and fusion of MVBs with the tonoplast is relatively well described and known to involve factors implicated in vacuolar maintenance^9,10,17^. Consequently, disruptions in functionality of MVBs greatly impact vacuolar morphology^9,10,18,19^. Importantly, cells with aberrant MVB machinery are not devoid of the tonoplastic material, but typically contain large amount thereof. However, these mutants fail to organize the tonoplast into an organelle with the typical vacuolar morphology^9,11,19^. Furthermore, some of the mutants with impaired MVB functionality have highly aberrant vacuolar morphology in young cells, where the organelle seems to undergo intensive fission and fusion, but relatively normal vacuolar morphology in the differentiated cells^9,19^. These data strongly indicate that MVBs play an important role in regulating the tonoplast organization but do not serve as the initial source of the vacuolar membrane.

Furthermore, a number of previous studies implementing whole-cell 3D TEM^20^, single-section TEM^2^ or confocal microscopy^3,21^ reported that plant vacuoles at the early stages of their establishment comprise a dynamic tubular network that might undergo temporary fragmentation during cell division. Cui and colleagues suggested that the tubular morphology of young vacuoles observed in the previous studies might be an artifact caused by fluorescent probes used to visualize the organelle, i.e. BCECF and SNARF^7^. Moreover, authors pointed out that the existing studies offer broad conclusions supported only by fragmented data obtained on specific developmental stages in different cell types. We endeavoured to address these issues and performed a series of experiments that provided a systematic overview of the vacuolar establishment process and allowed us to re-evaluate our previous findings.

We focused on implementing *Arabidopsis thaliana* primary root as a model, as it is an excellent system for tracking successive stages of cell differentiation and is thus optimal for studying vacuolar development. Root growth is a result of cell division and differentiation. The former occurs close to the root tip in the meristematic zone (MZ), which is maintained by the quiescent center (QC). Stem niche cells located in proximity of the QC, give rise to lineages of cell types that undergo gradual differentiation and elongation^22^ (**Fig. 1a**). As a result, a single vertical tier of cells will represent a snapshot of successive developmental stages, with the earliest steps occurring close to the QC and the latest taking place proximal to the hypocotyl.

**Figure 1.**
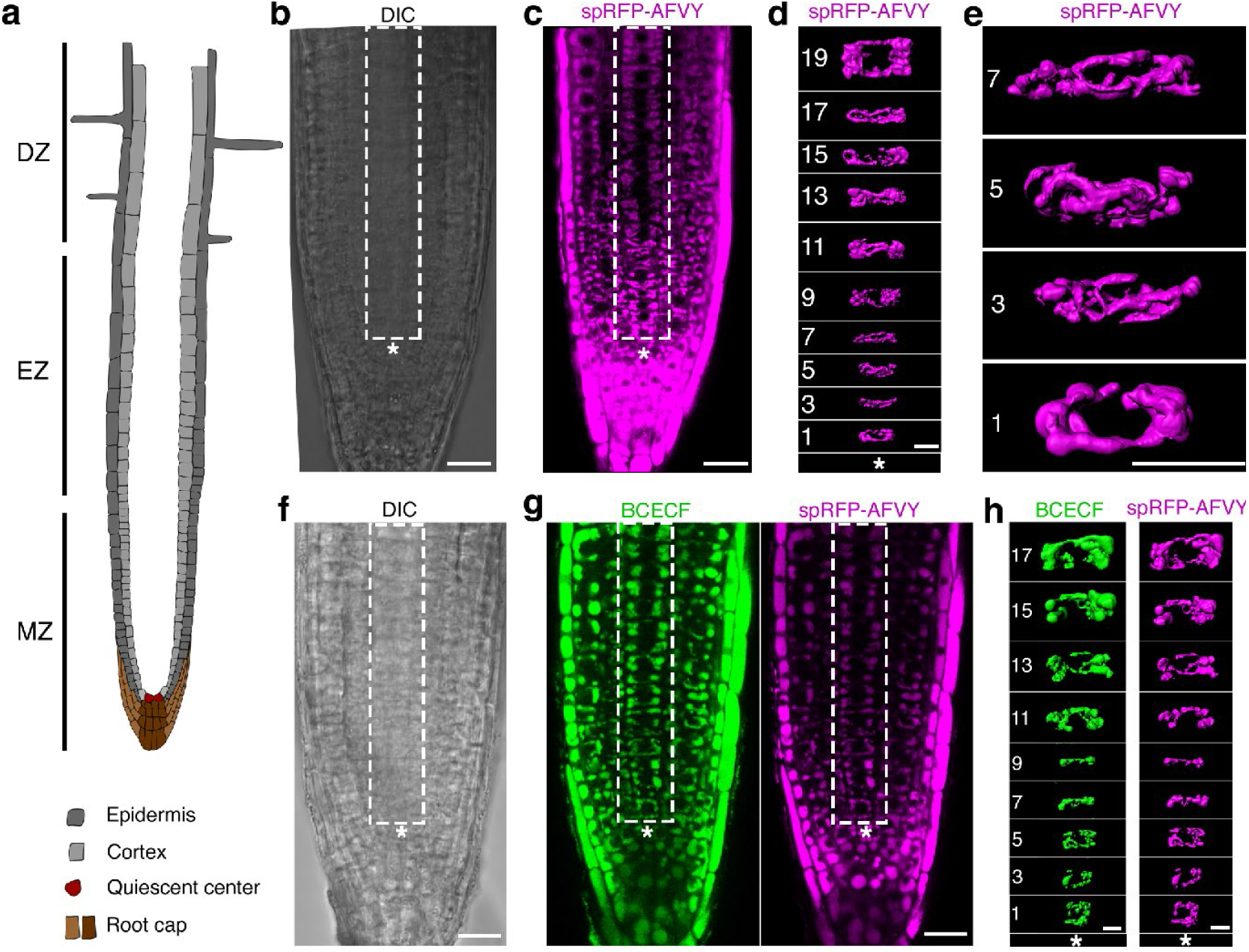
Vacuolar morphology at successive stages of biogenesis was visualized using fluorescent dye BCECF and endogenously expressed vacuolar marker spRFP-AFVY. **(a)** A scheme of Arabidopsis root depicting cell types presented in this study. MZ, meristematic zone containing youngest cells with the earliest stages of vacuolar biogenesis close to the QC. EZ, elongation zone contains cells that entered differentiation program resulting in their elongation along vertical axis of the root. DZ, differentiation zone is generally defined by the presence of highly elongated cells and root hairs. *Arabidopsis thaliana* seedlings expressing vacuolar marker spRFP-AFVY were either directly scanned using CLSM **(b-e)** or first stained with the BCECF dye typically used to visualize vacuolar lumen **(f-h)**. Obtained z-stack scans were then processed using Imaris software to produce 3D renderings of vacuoles in cortex cells of a single tier (rectangular selection). Successive stages of vacuolar development were tracked from the youngest cells of the root (closest to quiescent center (QC), asterisk)), cell position was numbered counting from the QC. Note, that staining with BCECF shows no discernible effect on vacuolar morphology. Importantly, vacuoles have a distinctly tubular morphology even in the youngest cells of the root cortex. **(b)** DIC channel, a single optical slice of a z-stack scan of an *Arabidopsis thaliana* root to show the position of the quiescent center (marked with an asterisk.) Rectangular selection denotes the area used for the 3D reconstruction shown in (**d**). **(c)** RFP channel, a single optical of the same z-stack as in (**b**), to show position of the cortex cells used for rendering in (**d**). (**d**) 3D rendering of the z-stack illustrated in (**b**) and (**c**) showing vacuoles at successive developmental stages in the single tier of cortex cells. (**e**) Enlarged view of cells presented in (**c**) showing vacuoles in the youngest cortex cells, numbers indicate cell position from the QC. **(f)** DIC channel, a single optical slice of a z-stack scan of an *Arabidopsis thaliana* root to show the position of the quiescent center (marked with an asterisk.) **(g)** Green (BCECF) and magenta (spRFP-AFVY) fluorescent channels, a single optical slice of a z-stack scan of an *Arabidopsis thaliana* root to show the position of the cortex cells used for rendering in (**h**). **(h)** 3D rendering of the z-stack illustrated in (**f**) and (**g**) showing vacuoles at successive developmental stages in the single file of cortex cells. *, QC. Numbers indicate cell position from the QC. White rectangles denote area positions used for 3D reconstruction of vacuoles in cortex cells. Scale bars: 10 um

In this study, we focused on implementing confocal laser scanning microscopy (CLSM,) which enables *in vivo* tracking of markers localized in plant vacuoles, thus allowing quantitative assessment of the content exchange between individual vacuolar substructures.

To assess if BCECF, a dye commonly used to visualize the vacuolar lumen^23^, affects morphology of the organelle, we compared phenotypes of vacuoles in *Arabidopsis thaliana* root cells in the absence or presence of the dye^23^ (**Figure 1b-h**). In these experiments we used two independent transgenic lines expressing red fluorescent protein delivered to the vacuole implementing N- or C-terminal propeptide sorting signals^24,25^, spL-RFP and spRFP-AFVY, respectively^26^. To ensure that our observations are representative of vacuolar biogenesis, we assessed vacuolar morphology at successive developmental stages of the same cell type. For this, we performed high-resolution 3D confocal microscopy of an area encompassing the root meristem and elongation zones. The imaging data was then used to select a single column of cortex cells, in which cell positions were numbered counting from the QC and 3D reconstructions were generated for the vacuoles of every second cell (**Figure 1b-e, Movie S1**). We selected cortex cells for analysis, since the 3D TEM reconstructions presented by Cui and co-authors were predominantly done on this cell type^16^.

Consistent with previously published data based on CLSM^3,7^, single section TEM^2^ and whole-cell 3D TEM^20^, we observed a fine tubular vacuolar network in young plant cells (**Figure 1e**). The diameter of the tubules was gradually increasing with the age of the cells, eventually causing fusion of the network into balloon-like vacuolar structures that occupied most of the volume in differentiated cells (**Figure 1b-e, Movies S1 and S2**). We then checked the effect of BCECF on the vacuolar morphology. To this end, seedlings of the same transgenic lines as described above were stained with BCECF and imaged using the same protocol. Our data unequivocally demonstrated that BCECF does not cause discernible changes in vacuolar morphology at any of the observed stages of vacuolar biogenesis (**Figure 1f-h**). Thus, we confirmed the validity of the results from the previous studies implementing BCECF staining.

Additionally, we performed time-lapse imaging of cortex cells at successive developmental stages to visualize the mobility of the vacuolar structures (**Movie S2**). The uninterrupted directionality of vacuolar structures movement in young cells strongly implied that they are connected. However, resolution of CLSM is not sufficient to conclusively discern whether we were observing movement of connected tubular vacuolar network or chains of tethered small vacuoles. To assess whether vacuolar structures indeed shared the lumenal space, we attempted to track diffusion of fluorescent markers in the vacuolar lumen using a FRAP (Fluorescence Recovery After Photobleaching) assay (**Figure 2a-d**). In brief, the fluorophore molecules within a small vacuolar area were exposed to a pulse of high intensity excitation light resulting in an irreversible loss of their fluorescence (photobleaching), after which re-distribution of the fluorescence intensity within the vacuolar area (fluorescence recovery) was tracked to assess diffusion of fluorophore molecules between the vacuolar substructures of the cell.

**Figure 2.**
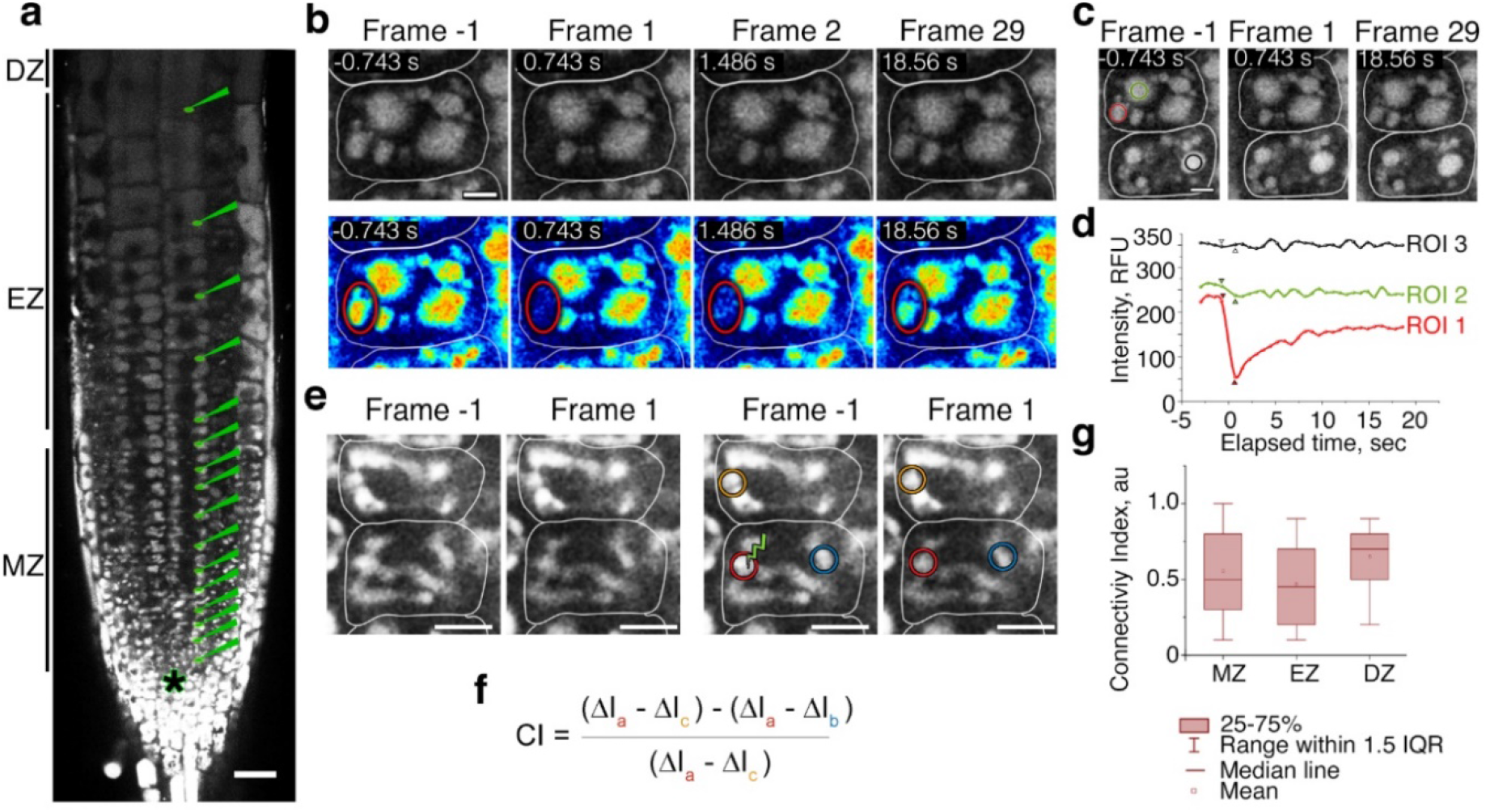
FRAP assay reveals that vacuolar structures represent a connected network even at the earliest stages of vacuolar development. *Arabidopsis thaliana* seedlings expressing vacuolar marker spL-RFP were subjected to FRAP assay. (**a**) The photobleaching was performed on every second cortex cell, counting from the QC. For each cell, a small portion of vacuolar area was subjected to photobleaching. The fluorescence recovery was then used to estimate whether vacuolar structures within each cell comprise a connected network. Scale bar, 10 um. *, QC. (**b**) Frames from a FRAP time-series obtained on a meristematic cell containing seemingly separated vacuolar structures. Top row shows the raw data, bottom row shows the same data colour-coded for signal intensity using Royal LUT. White lines denote cell walls, the red oval indicates a vacuolar structure within which a small spot was exposed to high intensity laser during photobleaching. Four frames were acquired prior to and 25 frames were acquired after the photobleaching. Scanning was performed at the speed of one frame per 743 msec. “Frame −1”, the frame acquired right before photobleaching. “Frame 1”, the first frame after photobleaching. Time in seconds, elapsed time from the photobleaching event. Increase in fluorescence within photobleached vacuolar structure is visible already on the Frame 2, less than two second after photobleaching occurred. Such rapid dynamics of recovery strongly indicates free diffusion of the fluorophore between vacuolar structures visible on the optical section of the cell. Scale bar: 5 um. (**c**) Areas used for quantification of the FRAP imaging data illustrated in (**b**). Regions of interest (ROIs) were selected to represent the photobleached region of the vacuole (red), a vacuolar structure inside the same cell, which was not exposed to high intensity laser light (green) and a vacuolar region in a neighbouring cell (blue). (**d**) The change of fluorescence intensities inside ROIs shown on (**c**) were measured for the FRAP time-lapse data. The arrows indicate data points corresponding to the Frame −1 and Frame 1. ROI1 and ROI2 show decrease in the intensity right after photobleaching, indicating that they are located in connected structures. Expectedly, the ROI3, which corresponds to a vacuolar area of another cell, does not show decrease in the intensity. Half time of fluorescence recovery in the photobleached area is shorter than 2 seconds, consistent with free diffusion of the fluorophore into the photobleached area. (**e**) Frames from a FRAP series performed on highly tubulated vacuoles of young cortex cells. “Frame −1”, the frame acquired right before photobleaching. “Frame 1”, the first frame after photobleaching. White lines indicate cell walls. Scale bar: 10 um. The left panel shows the raw data, the right panel shows the same data with indicated areas used for Connectivity index calculation. (**f**) Equation for the Connectivity index (CI) calculation, colored letters a, b, c indicate areas denoted on (**e**) with the corresponding colors. CI = 1 indicates that photobleached and not photobleached areas are fully connected and diffusion rate between them is higher than the scanning speed. CI values higher than 0.1 indicate localization of the fluorophore in connected compartments, CI values lower than 0.1, indicate that fluorophore is trapped in separated compartments. For more information see Figure S1. (**g**) A two-tailed Students t-test comparing CI indexes calculated for the cells belonging to the MZ, meristematic zone (cells at the positions 1-13, counted from QC); EZ, elongation zone (cells at the positions 14-19), DZ, differentiation zone (cells at the positions 20 and upwards) revealed similar connection of the vacuolar structures in the cells of all three zones (p-value > 0.05). The chart shows representative data of one out of six FRAP experiments, that comprised use of 7 roots to acquire 63 FRAP series.

In line with our 3D reconstruction approach described above, we made sure to provide systematic data and performed photobleaching at successive developmental stages in the cortex cells counting their position from the QC. FRAP assay of spL-RFP and spRFP-AFVY lumenal markers produced similar results (**Figure 2, Movie S3**), i.e. at all developmental stages fluorescence recovery was observed within a couple of seconds, consistent with diffusion of the marker inside a connected vacuolar network (**Movie S3)**. Even in cells containing seemingly fragmented vacuoles, we observed a rapid recovery of fluorescence in a photobleached vacuolar portion, proving that also these cells contain vacuolar networks (**Figure 2b-d**).

Performing FRAP assays on young plant vacuoles is a notoriously challenging task. The small size of the young vacuolar structures and their rapid movement during time-lapse imaging causes fluctuations in the fluorescence intensity that interferes with accurate quantification of the intensity changes originating from the diffusion of the fluorophore. Thus applying the traditional FRAP quantification methods to the highly mobile vacuolar networks of the youngest meristematic cells was yielding unreliable data. We overcame this issue by developing a customized approach for quantifying FRAP data to calculate the connectivity index (CI) of plant vacuoles (**Figure 2e-g, Figure S1,** for the detailed description see the Methods chapter). CI value of 1 indicates unrestricted diffusion of a fluorophore inside the vacuolar lumen and CI values below 0.1 would represent a lack of diffusion due to the fluorophore localization in separated compartments. We calculated connectivity indexes for cells of the meristematic zone (MZ) which contain the youngest vacuoles, elongation zone (EZ) comprising cells with vacuoles that underwent substantial fusion and differentiation zone (DZ) in which cells contain large vacuoles connected into balloon-like structures. Statistical analysis revealed no significant difference in connectivity indexes for all three zones (**Figure 2g**), thus confirming that even at the earliest stages of their development plant vacuoles comprise a dynamic network of connected tubules rather than separate vesicles.

Notably, Cui and colleagues observed fragmentation of vacuoles not only while using super-resolution microscopy on the fixed cells, but also in the experiments involving photoconversion of a vacuolar marker in the living cells. It did not escape our attention that their photoconversion experiments revealed fragmented vacuoles in elongated cells (cell length: width more than 2:1). Which should in wild-type plants under normal growth conditions typically contain connected vacuoles.

It is well established that depending on environmental stimuli, plant vacuoles can undergo drastic morphological changes, including fragmentation^1,3,4,16,27–29^. Therefore, we speculated that the discrepancies between our observations and the data reported by Cui and colleagues might originate from differences in plant growth conditions. To test reproducibility of our results, we compared the morphology of vacuoles in seedlings grown in two independent laboratories in two different countries. Eight independent experiments performed at two different research facilities consistently revealed a tubular vacuolar network in young root cells (**Figure S2**). Therefore, we are confident in our conclusion that under growth conditions considered as standard by the plant research community, young plant vacuoles comprise a connected tubular network. Thus, our results provide a systematic evidence that validates, corroborates and unifies previous observations^3,7,2,20^.

In summary, in this study: (i) we argue that the existing knowledge on plant endomembrane trafficking does not support the hypothesis of vacuolar structures originating from homotypic fusion of MVBs; (ii) we provide a comprehensive overview of plant vacuolar morphology at successive developmental stages starting from the youngest cells proximal to the QC and demonstrate that even at the earliest stages vacuoles have a tubular network morphology; (iii) we determine that BCECF does not affect vacuolar morphology, thus validating conclusions from previous studies using this dye; (iv) we use a customized FRAP assay to provide a systematic assessment showing that plant vacuoles comprise a connected network already at the earliest stages of development.

## Methods

### Plant material and growth

In this study we used Arabidopsis thaliana Col-0 accession wild-type plants and transgenic plants expressing spL-RFP and spRFP-AFVY markers^26^. Seeds were surface sterilized for 20 min in 70% Ethanol containing 0.05% Triton X-100, washed in 95% Ethanol and air dried. Seedlings were grown on vertically positioned plates containing 0.5x MS (M0222, Duchefa), supplemented with 10 mM MES (M1503, Duchefa), 1% sucrose and 0.8 % Plant agar (P1001, Duchefa), pH5.8, under long day conditions (150 uM light, 16h light, 8h darkness, 22°C)

### BCECF staining

BCECF-AM (2’,7’-Bis-(2-Carboxyethyl)-5-(and-6)-Carboxyfluorescein, Acetoxymethyl Ester, B1150 ThermoFisher) was stored as a 10 mM stock solution in DMSO. For staining, 5-7 days old seedlings were submerged in liquid 0.5xMS (0.5x MS (M0222, Duchefa), 10 mM MES (M1503, Duchefa), 1% sucrose pH5.8) containing 10 μM BCECF and incubated at room temperature for 2h, followed by washing for 15 min in fresh 0.5xMS liquid medium and immediate mounting for imaging.

### Confocal microscopy

Images were obtained using Leica SP8 and Leica SP5 confocal microscopes with LAS X software and Zeiss LSM 800 with ZEN software. To avoid possible dehydration of samples during prolonged scans, seedlings were mounted in 0.5x liquid MS in a RoPod 1 chamber (https://www.alyonaminina.org/ropod), roots were covered with a coverslip and the chamber lid was closed to maintain humidity during imaging.

### 3D

Scanning was performed using Leica SP8 CLSM with 63x objective, NA 1.30. BCECF was imaged using 488 nm excitation light, 490nm - 552nm emission range and the standard mode of the HyD detector. RFP fluorescence was excited using 561 nm light and 577nm - 700nm emission range was detected using the standard mode of the HyD detector. Pinhole was set to 1AU, scanning speed was set to 400 Hz, line average to 3, pixel resolution was set to optimal and number of optical slices was system optimized.

Most of the experiments were performed at COS, Heidelberg University, Germany. Additionally, observed vacuolar phenotypes were tested on seedlings grown at Uppsala BioCenter, SLU, Sweden using a Zeiss LSM 800 microscope equipped with GaAsp detectors and ZEN 2 software, 63x water immersion objective, NA 1.20. Excitation and emission settings were set similarly to the ones described above.

### 3D reconstruction

Imaris 9.2 (Bitplane) was used for 3D reconstructions of vacuoles. For each reconstructed vacuole, cell walls of a corresponding cell were manually traced using at least every third slice of the z-stack based on the DIC channel. The tracing data was then used to create a cell surface reconstruction. In the next step, the surface of the analysed cells was used to create a masked channel, in which signal intensity for the voxels outside the selected cell surface were set to 0. This channel was then employed to automatically create a 3D model of vacuolar lumen of the analysed cell. The obtained model was visually compared and adjusted to the underlying signal with the background subtraction threshold option. Reconstructions of vacuolar network for individual cells were then combined into a common overview image shown on the (**Figure 1d, h**).

### 4D

Time-lapse imaging was performed with the same settings for excitation and emission, as used for 3D imaging. The scan area size was cropped to enable frame rate of approximately one frame per second.

### FRAP

Imaging for FRAP assay was performed using Leica SP5, 40.0x objective NA 1.20. The excitation and emission settings were the same as used for obtaining 3D scans. To reduce photobleaching caused by scanning and speed up frame acquisition rate, resolution was set to 512×512 pixels, scanning speed to 600 Hz and line averaging to 1. FRAP settings were adjusted using FRAP tool of LAS AF software. A crosshair tool was used to select a single dot inside vacuolar area to be exposed to high intensity laser. Photobleaching was performed for 1 second using laser intensity empirically adjusted to achieve at least 30% drop in fluorescence intensity in the selected spot. Four frames were acquired prior to photobleaching to obtain data on the normal fluctuations in the marker fluorescence intensity caused by scanning. 25 more frames were acquired after photobleaching to track signal recovery. Each FRAP time-series was ca 21 seconds in length with approximately 740 msec/frame.

FRAP quantification was performed using a custom designed macro for automated extraction of FRAP series from Leica project file and a slightly modified FRAP profiler plugin for ImageJ (https://github.com/AlyonaMinina/vacFRAP). The adjustments of the plugin included hardcoding presence in the time-lapse of 4 frames prior to photobleaching and thus building the recovery curve fit starting from the 5^th^ frame.

### Connectivity index

High motility of young vacuoles was causing significant fluctuations of fluorescence intensity within a selected area and masking intensity changes originating from the diffusion of the fluorophore. Thus standard FRAP approach could not be efficiently applied for our samples. Hence we developed a customized approach. We argued that diffusion of the fluorophore within aqueous solution of a vacuole is a rapid process and photobleaching of a small area would lead to a detectable drop of fluorescence in the whole vacuolar network. To assess the rapid changes in the fluorophore distribution we analysed FRAP series frames right before (Frame −1) and right after (Frame 1) photobleaching, measuring changes in the fluorescence intensities within selected areas. On these frames we selected a small vacuolar area that was exposed to high intensity laser (area “a” within the red circle on the **Figure S1**) and an area on of the vacuolar network most distant from the photobleached region (area “b” within the green circle on the **Figure S1**). Furthermore, an area in the vacuolar network of neighbouring cell (area “c” within the blue circle on the **Figure S1**) was used as a control, since its vacuole was definitely not connected to the vacuolar network of the assessed cell.

Connectivity index was then calculated in following steps:

- ΔIa = (Ia_(Frame 1)_ - Ia_(Frame −1)_)*100/ Ia_(Frame 1)_; intensity drop in the photobleached area of the vacuole as % of this area intensity before photobleaching
- ΔIb = (Ib_(Frame 1)_ - Ib_(Frame −1)_)*100/ Ib_(Frame 1)_; intensity drop in the not photobleached area of the same vacuole as % of this area intensity before photobleaching
- ΔIc = (Ic_(Frame 1)_ - Ic_(Frame −1)_)*100/ Ia_(Frame 1)_; = intensity drop in the vacuole of neighbouring cell as % of this area intensity before photobleaching
- Rel = ΔIa-ΔIb; relative loss of fluorescence within the vacuole, comparing photobleached and non-photobleached areas (areas are in potentially connected parts of vacuole) Ref = ΔIa - ΔIc;reference loss of fluorescence, comparing photobleached area in one cell and non-photobleached in another (areas are in disconnected vacuoles) CI = (Ref-Rel)/Ref; Connectivity Index

CI = 0 indicates that photobleached and not photobleached areas are not connected. CI = 1 indicates that photobleached and not photobleached areas are fully connected and diffusion rate between them is higher than the scanning speed. Cut-off value, the lowest CI value that corresponds to connected network was estimated empirically. To establish the cut-off value for this study we used cells of the differentiation zone as the positive control representing connected vacuoles and vacuoles in neighbouring cells as a negative control, representing disconnected vacuolar structures (**Figure 2a** and **Figure S1**). Under our conditions, the CI values lower than 0.1 reproducibly correlated with not connected vacuolar compartments. Please note, that CI <0 means that non-photobleached area lost more signal than the photobleached area, indicating technical issues most probably resulting from sample drift during scanning. Similarly, CI > 1 indicates that there is a technical issue, most probably sample drift during scanning.

Connectivity indexes were quantified using the designated semi-automated assay implementing ImageJ macros (https://github.com/AlyonaMinina/Connectivity-Index). The macros were designed to first process Leica project files and extract individual FRAP series as tiff stack files and then guide the user for selecting the areas required for quantification of the CI index.

## Supporting information

Movie S1. 3D reconstruction of plant vacuoles in the root cortex cells at successive developmental stages reveals tubular network morphology.

Movie S2. Live imaging of plant vacuoles at successive stages of development.

Movie S3. FRAP at successive stages of vacuolar development.

Figure S1. Connectivity Index allows quantitative comparison of fluorescence recovery in highly mobile vacuolar structures of young root cells

Figure S2. Tubular vacuoles are reproducibly detected in the young cortex cells of seedlings handled at two independent research facilities

## Acknowledgements

This study was supported by Marie Sklodowska-Curie Actions Individual Fellowship (MAPoPHAGY, 799433) to E.A.M. and by the German Research Foundation (DFG; SCHE 1836/4-1) to D.S.

Authors would like to express their deep gratitude to Prof. Guido Grossmann for the intellectually stimulating coffee-break discussions that greatly benefited this study.

## Competing interests

Authors declare no competing interests.

---

**Movie S1.**
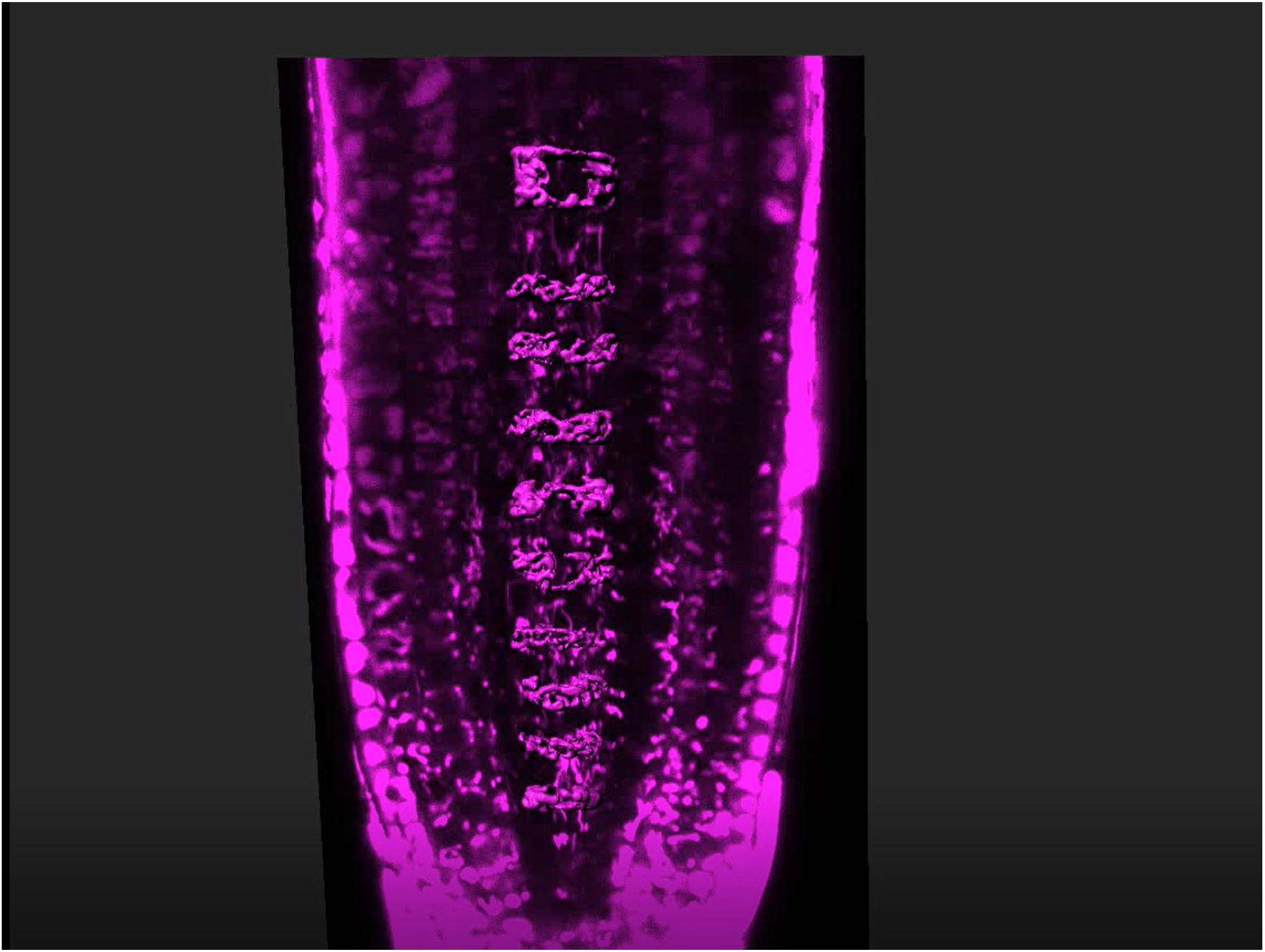
3D reconstruction of plant vacuoles in the root cortex cells at successive developmental stages reveals tubular network morphology. spRFP-AFVY, red fluorescent marker for vacuolar lumen, was detected in *Arabidopsis thaliana* roots using CLSM. A single tier of cortex cells was selected to reconstruct vacuolar structures at successive stages of cell differentiation. The imaging and reconstruction were performed as described in the Materials and methods chapter.

**Movie S2.**
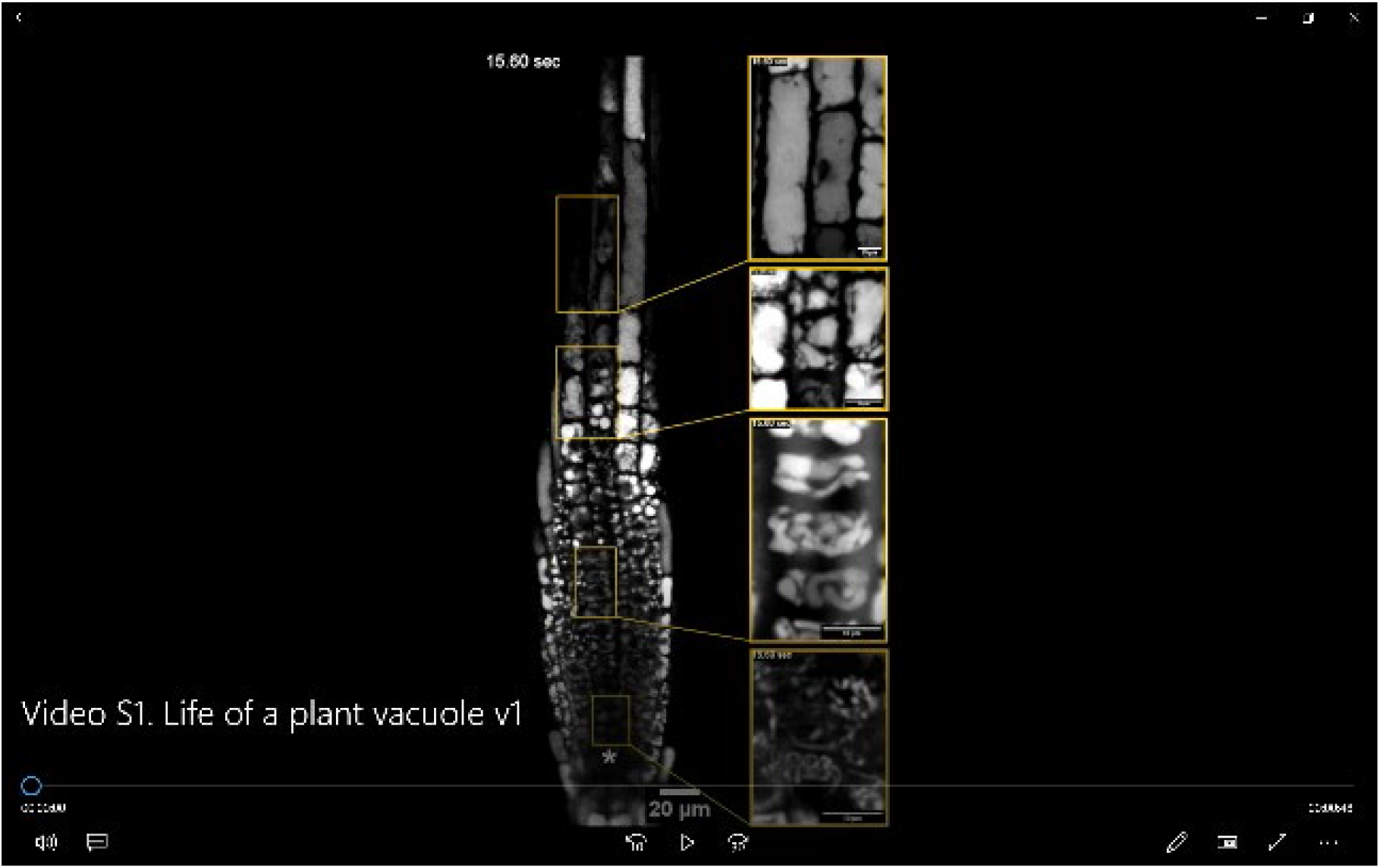
Live imaging of plant vacuoles at successive stages of development. *Arabidopsis thaliana* roots expressing spL-RFP, red fluorescent marker for vacuolar lumen, were imaged using CLSM. Time-lapse images of a single optical section show highly dynamic tubular vacuoles in young cells of the cortex. During cell differentiation and elongation, the thin vacuolar tubules gradually swell and eventually fuse into balloon-like structures. *, Quiescent Center; scale bar: 10 um.

**Movie S3.**
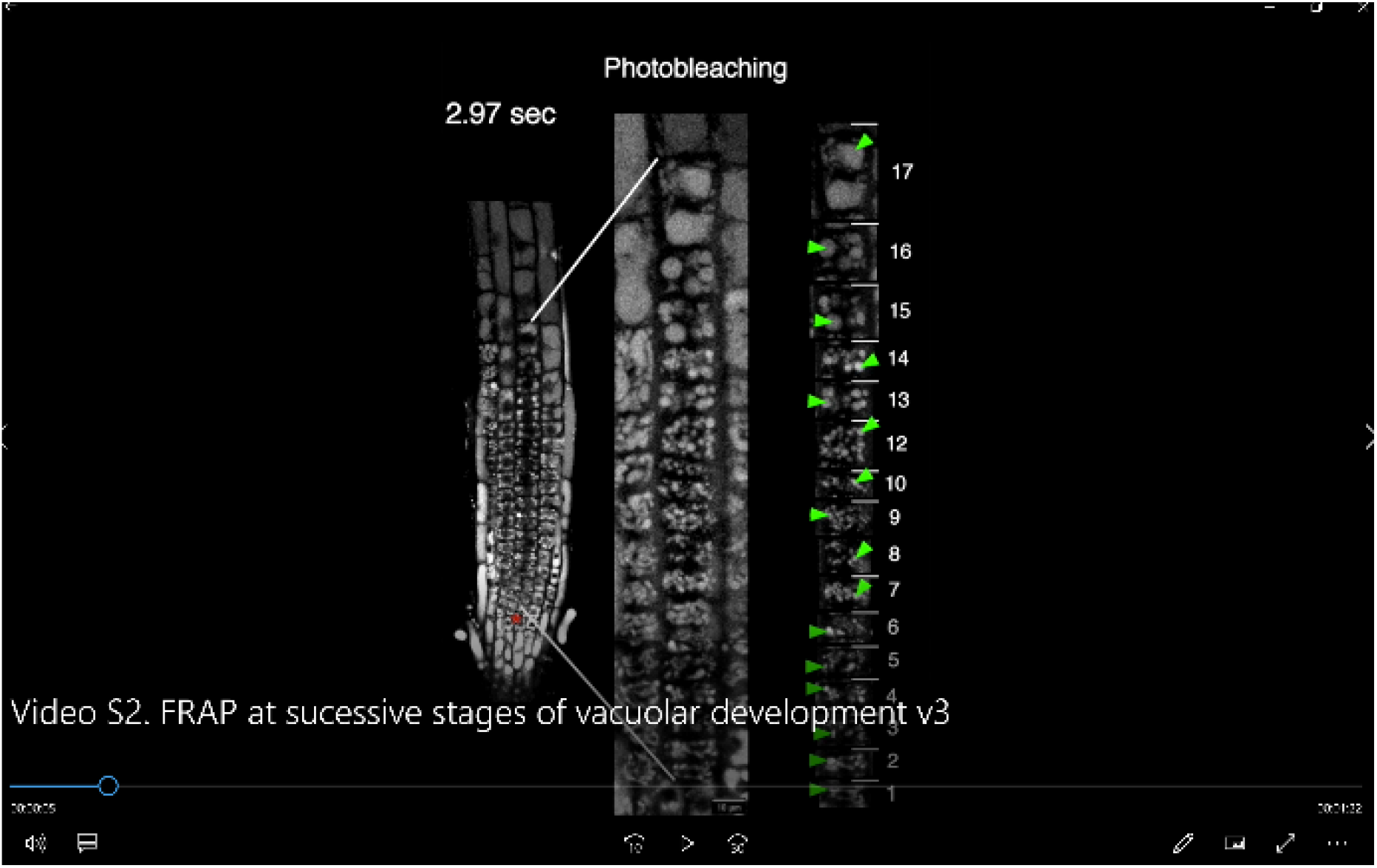
FRAP at successive stages of vacuolar development. *Arabidopsis thaliana* seedlings expressing spRFP-AFVY, a fluorescent marker for vacuolar lumen, were subjected to FRAP assay. For each cell, 4 scans were acquired prior to photobleaching, recovery after photobleaching was tracked on 25 subsequent scans. The overview of the root on the left shows the position of the quiescent center (QC, asterisk) and position of the cortex cells tier used for the analysis. Numbers indicate the cells position when counted from the QC. FRAP was performed on 17 consecutive stages of vacuolar establishment to systematically assess possible changes in the vacuolar connectivity during their establishment. At all observed stages we detected fluorescence recovery occurring within the few seconds, consistent with free diffusion of the fluorophore in the connected volumes of the vacuolar network. Quantitative comparison of vacuolar connectivity in the meristematic, elongation and differentiation zones is shown on the **Fig. 2g**. Scale bar: 10 um.

**Figure S1.**
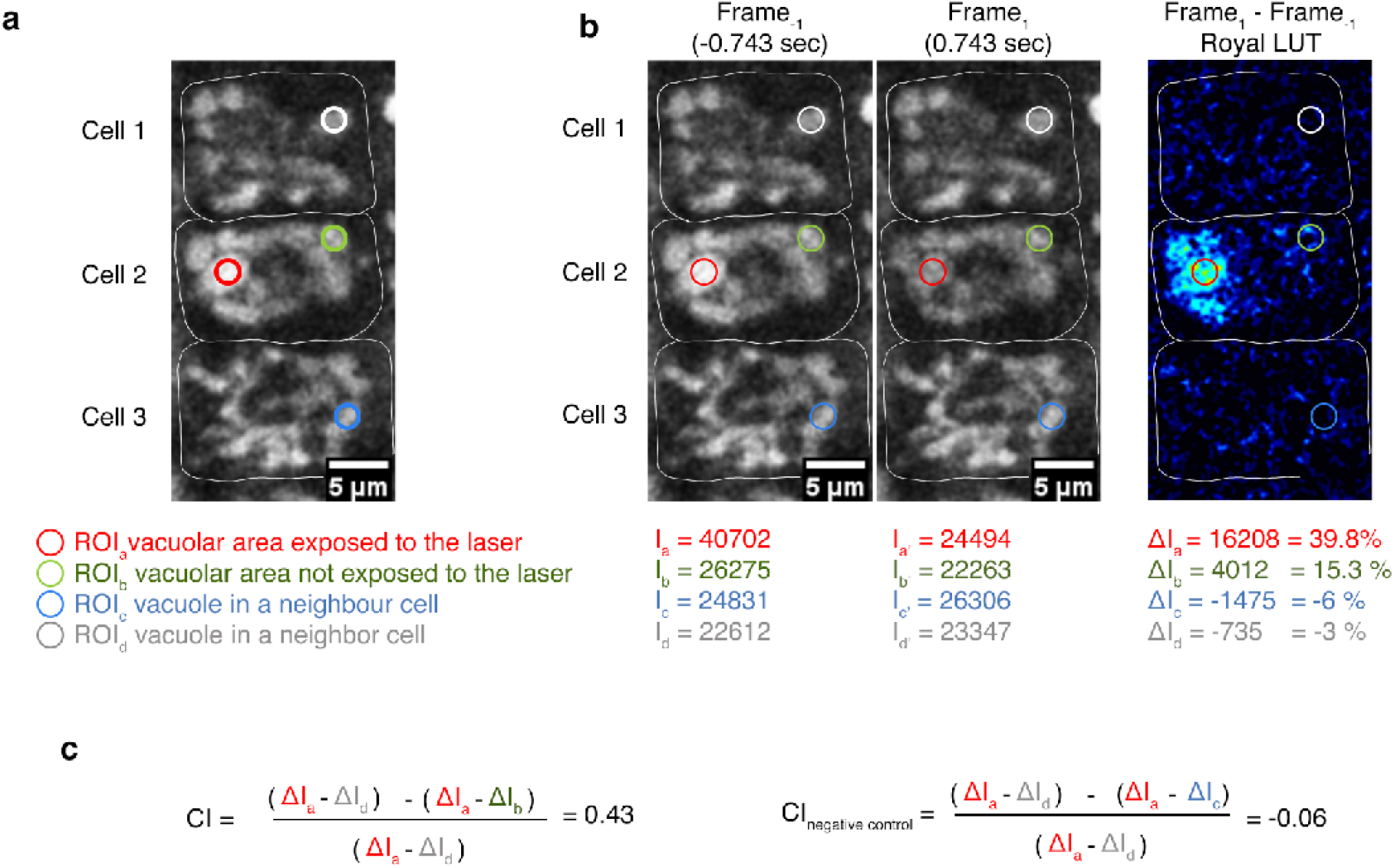
Connectivity Index allows quantitative comparison of fluorescence recovery in highly mobile vacuolar structures of young root cells. (**a**) FRAP assay was performed according to the description provided in the Materials and methods chapter using two independent transgenic Arabidopsis lines expressing red fluorescent markers for vacuolar lumen. An example of regions or interest (ROIs) selected on 3 cells, among which only the ROI1 in the cell 2 was exposed to high intensity laser. White lines indicate cell walls. (**b**) Frame_-1_ was taken immediately before the start of photobleaching. Frame_1_ was acquired right after photobleaching was finished. The numbers signify fluorescence intensity values within the regions of interest measured using ImageJ software. To highlight the changes in the intensity caused by photobleaching, Frame_1_ image was subtracted from Frame_-1_ image using ImageJ “Calculator” function. The resulting image was color-coded using Royal LUT, lighter colors indicate bigger difference in the intensity. Numbers show fluorescence intensity decrease within the ROIs, which are presented as percent of the fluorescence signal within the corresponding ROI before photobleaching. (**c**) Connectivity Index (CI) is calculated by comparing change in fluorescence in ROIs within the same cell and in neighbouring cells. Comparison of ROIs within the same cell reveals diffusion rate of the fluorophore within vacuolar structures of the cell. Comparison of ROIs located in different cells informs about fluctuations in fluorescence intensity caused by scanning and movement of the vacuoles. CI values higher than 0.1 indicate that fluorophore can diffuse between vacuolar structures of the cell. CI values lower than 0.1 indicate that the fluorophore is trapped within separated structures.

**Figure S2.**
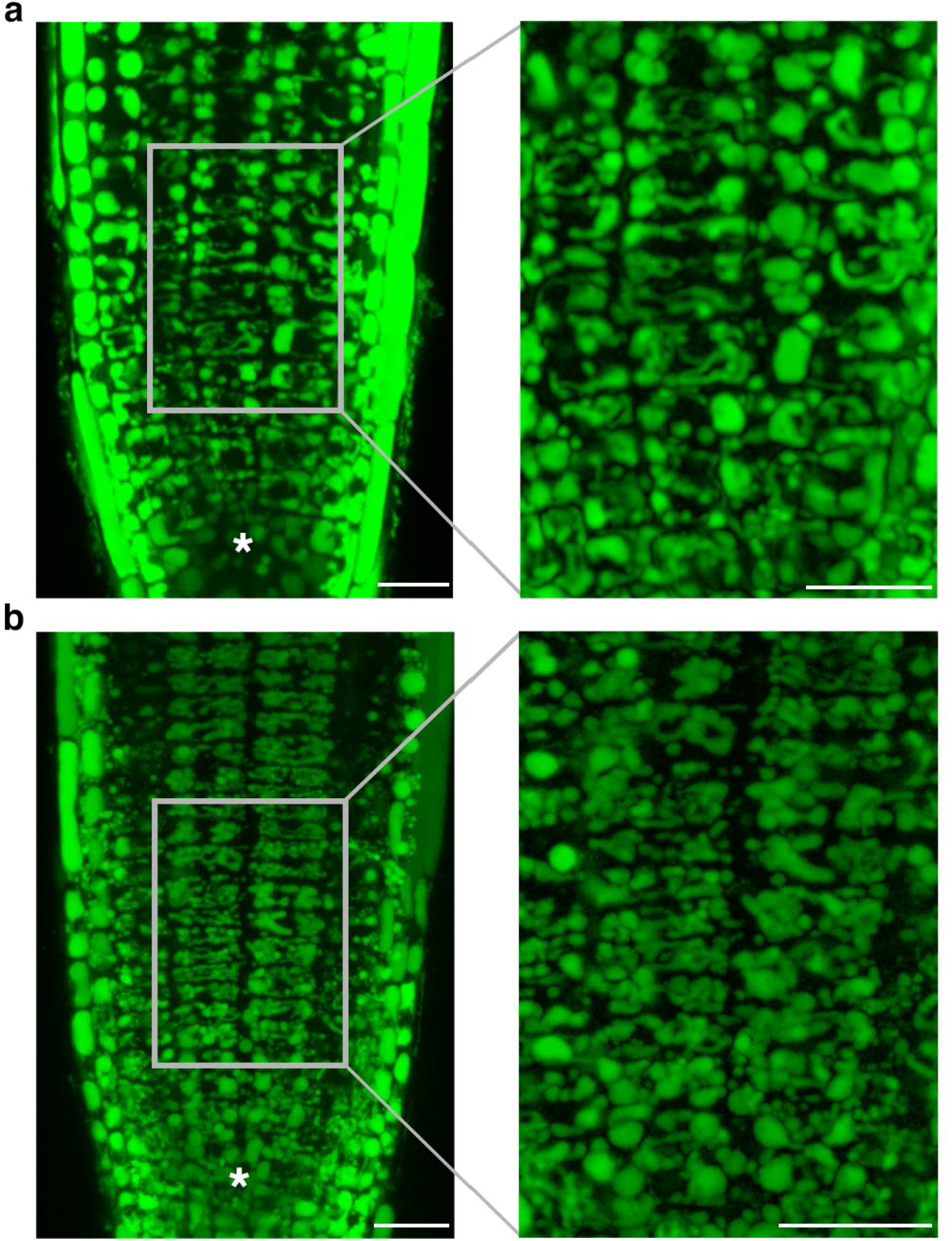
Tubular vacuoles are reproducibly detected in the young cortex cells of seedlings handled at two independent research facilities. (**a**) spRFP-AFVY transgenic line was grown in the COS plant growth facility, University of Heidelberg, Germany. Five days old seedlings were stained with BCECF and imaged using Leica SP8 confocal microscope. Maximal projection of the BCECF channel for the part of the z-stack shows intricate tubular network in the young cells proximal to the quiescent center. (**b**) Col-0 wild type seedlings were grown in the plant facility of Uppsala BioCenter, SLU, Sweden. Five days old seedlings were stained with BCECF and imaged using Zeiss LSM 800 confocal microscope. Maximal projection of the BCECF channel shows young cortex cells containing tubular vacuolar network similar to the one shown on (**a**). *, QC. Scale bar: 20 um.

## Abbreviations

CLSM: confocal laser scanning microscopy
CTPP: C-terminal propeptide
BCECF, BCECF-AM: 2’,7’-Bis-(2-Carboxyethyl)-5-(and-6)-Carboxyfluorescein, Acetoxymethyl Ester
DZ: differentiation zone
ER: endoplasmic reticulum
EZ: elongation zone
FRAP: fluorescence recovery after photobleaching
ILVs: intraluminal vesicles
MVBs: multi vesicular bodies
MZ: meristematic zone
NTPP: N-terminal propeptide
QC: quiescent center
SVs: small vacuoles
TEM: transmission electron microscopy

## Notes

https://github.com/AlyonaMinina/Connectivity-Index

https://www.alyonaminina.org/ropod

### Competing Interest Statement

The authors have declared no competing interest.

### Summary of Updates

The main text and figure legends have been edited to clarify the message.

