## Supplementary figures and images for "Light at the end of the tunnel: FRAP assay reveals that plant vacuoles start as a tubular network"

### Figure S1. Connectivity Index allows quantitative comparison of fluorescence recovery in highly mobile vacuolar structures of young root cells

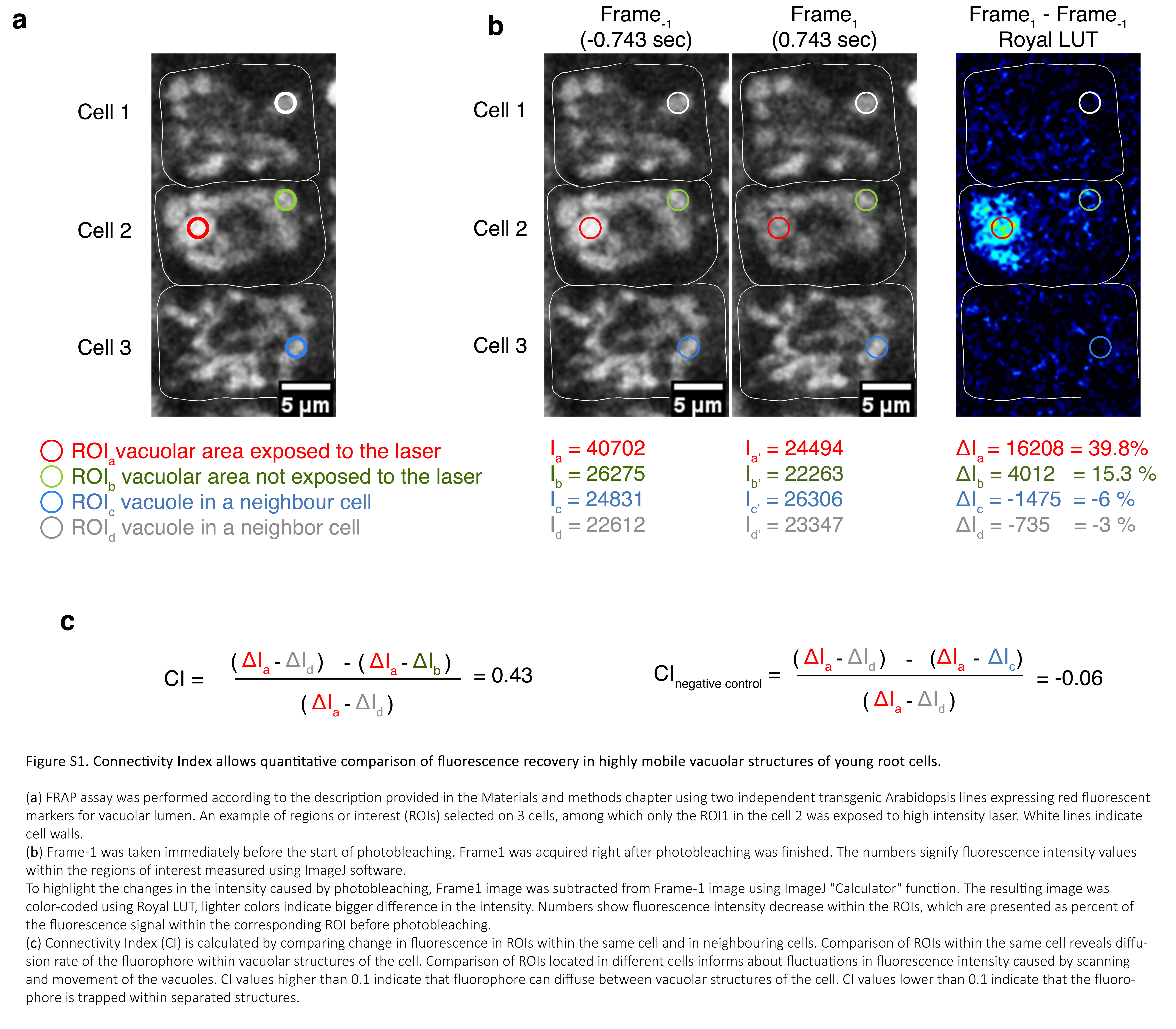

### Figure S2. Tubular vacuoles are reproducibly detected in the young cortex cells of seedlings handled at two independent research facilities

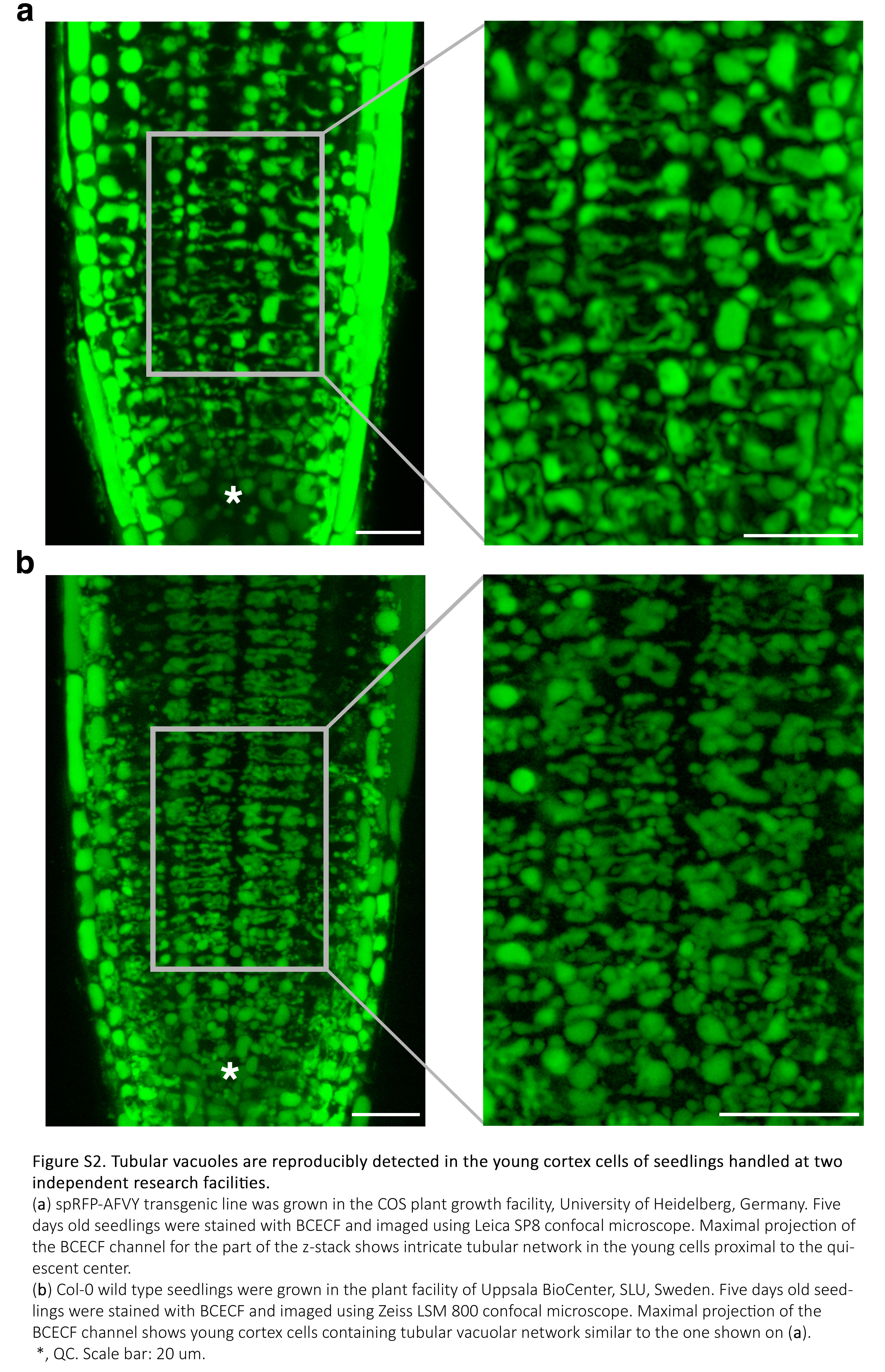
